# Efficient transgene-free multiplexed germline editing via viral delivery of an engineered TnpB

**DOI:** 10.64898/2026.01.23.700382

**Authors:** Trevor Weiss, Maris Kamalu, Honglue Shi, Gabriel Wirnowski, Alice Ingelsson, Jasmine Amerasekera, Kamakshi Vohra, Marena I. Trinidad, Zheng Li, Emily Freitas, Noah Steinmetz, Charlie Ambrose, Kerry Chen, Jennifer A. Doudna, Steven E. Jacobsen

## Abstract

Virus-induced genome editing (VIGE) using compact RNA-guided endonucleases is a transformational new approach in plant biotechnology, enabling tissue-culture-independent and transgene-free genome editing (Hu et al. 2025; Liu et al. 2025; Weiss et al. 2025). We recently established a VIGE approach for heritable editing at single loci in *Arabidopsis* by delivering the compact genome editor ISYmu1 TnpB (Ymu1) and its guide RNA (gRNA) via Tobacco Rattle Virus (TRV) (Weiss et al. 2025). Here, we greatly improved this approach by devising a multiple gRNA expression system and by utilizing an engineered high-activity Ymu1 variant (Ymu1-WFR) (Zhou et al. 2026) to develop an efficient multiplexed genome editing platform.

## Main Text

TRV is a bipartite RNA virus composed of RNA1 and RNA2. To evaluate TRV-mediated multiplexing capabilities, we co-delivered RNA1 with two RNA2 vectors encoding either *AtPDS3* gRNA12 or *AtCHLl1* gRNA4 to *Arabidopsis* (Figure S1A), two gRNAs with high activity (Weiss et al. 2025). Amplicon sequencing (amp-seq) revealed editing almost exclusively at one target site or the other (Figure S1B), suggesting viral superinfection exclusion (Perdoncini Carvalho et al. 2022). We therefore sought to develop a system in which both gRNAs could be expressed on a single RNA2 vector.

First, to find the optimal gRNA structure we identified the precise omega RNA (ωRNA) sequence via small RNA sequencing (RNA-seq) in *E. coli*, and found it to be 127-nucleotides (nt) in length (Figures S2A, S2B). In addition, we tested various ωRNA lengths and found that 127-nt gave the highest editing in protoplasts (Figures S2C, S2D). Using the 127-nt ωRNA, we tested multiplexed arrays featuring tRNA, HDV, HDV-HH, or a repeat as gRNA processing elements using an *Arabidopsis* protoplast assay (Figure S2F). Amp-seq analysis showed that while all designs enabled editing, HDV-based designs performed best at simultaneously editing both sites in protoplasts (Figure S2G). Furthermore, polymerase chain reaction (PCR) using primers spanning both sites suggested the occurrence of large deletions between the two target sites (Figure S2H). These experiments identified the HDV and HDV-HH designs as the top performing multiplexing arrays, and that Ymu1 TnpB can generate large deletions between two targeted sites.

To minimize RNA2 cargo size, the HDV multiplex array was selected for TRV-mediated multiplexed editing *in planta*. Initially, we designed two RNA2 vectors targeting *AtCHLl1* (gRNA4) and *AtPDS3* (gRNA12), incorporating a tRNA^Ileu^ mobility sequence at the 3’ end of the cargo to facilitate systemic movement and heritability (Figure S3A). gRNA4 targets the gene body of *AtCHLl1* (biallelic edits create yellow tissue sectors) and gRNA12 targets the promoter region upstream of the *AtPDS3* transcription start site (no visible phenotype). After delivering TRV vectors, we did not observe any phenotypic evidence of editing. Suspecting inefficient mobility or processing of the RNA2 cargo, we tested three additional constructs containing a tRNA^Ileu^ downstream of each HDV ribozyme (Figure 1A). After TRV delivery, yellow sectors appeared on leaves for all three vectors, indicating biallelic edits at *AtCHLl1* (Figure 1B). Yellow sectored plants infected with vectors pTW2278 and pTW2279 displayed average editing efficiencies of 25.6% for *AtCHLl1* and 30.2% for *AtPDS3* (Figure 1C), and those infected with pTW2498 showed 31.3% for *AtCHLl1* gRNA4 and 5.2% for *AtCHLl1* gRNA6 (Figure 1D).

**Figure 1:**
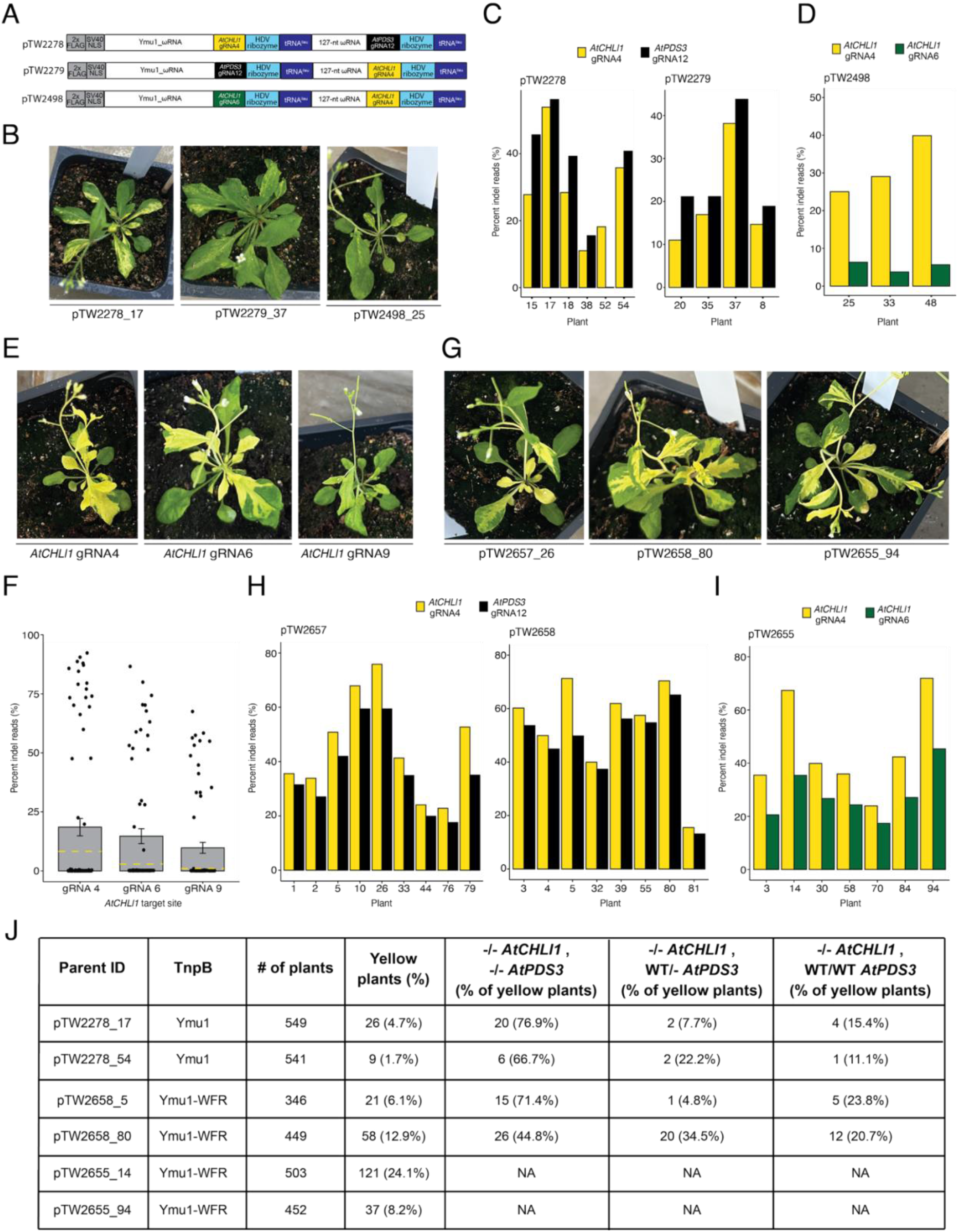
Heritable and transgene-free multiplexed genome editing in *Arabidopsis* via viral delivery of Ymu1. **(A)** Schematic representation of the cargo being expressed by the PEBV promoter in TRV RNA2. *AtCHLl1* gRNA4 (yellow), *AtPDS3 gRNA12* (black), and *AtCHLl1* gRNA6 (green) were used in the multiplex TRV experiments. The plasmid ID is listed to the left of each construct. **(B)** Yellow sector phenotype observed from plants infected with the TRV vectors from panel A. The plant ID (construct_plant) is listed below each picture. **(C and D)** Editing efficiency (y-axis) of plants displaying the yellow sector phenotype (x-axis) infected with TRV vectors pTW2278, pTW2279 and pTW2498. **(E)** Yellow sector phenotype observed from plants infected with TRV vectors expressing Ymu1-WFR targeting *AtCHLl1* (gRNA4, gRNA6, and gRNA9). The site being targeted is listed below each picture. **(F)** Editing efficiency (y-axis) of plants infected with TRV vectors expressing Ymu1-WFR targeting *AtCHLl1* (x-axis). Each dot represents an individual plant. The yellow dashed line on each bar indicates the average editing efficiency previously reported (Weiss et al. 2025). **(G)** Yellow sector phenotype observed from plants infected with TRV vectors expressing Ymu1-WFR targeting *AtCHLl1* gRNA4 and *AtPDS3* gRNA12 (pTW2657 and pTW2658) or *AtCHLl1* gRNA4 and *AtCHLl1* gRNA6 (pTW2655). The plant ID (construct_plant) is listed below each picture. **(H and I)** Editing efficiency (y-axis) of plants displaying the yellow sector phenotype (x-axis) infected with TRV vectors pTW2657, pTW2658 and pTW2655. (**J**) Heritability data from plant ID pTW2278_17, pTW2278_54, pTW2658_5, pTW2658_80, pTW2655_14, and pTW2655_94.

We recently engineered a highly active Ymu1 variant (Ymu1-WFR) (Zhou et al. 2026). To evaluate Ymu1-WFR efficiency using TRV, we targeted three published *AtCHLl1* target sites: gRNA4, gRNA6, and gRNA9 (Weiss et al. 2025). Infected plants showed a strong yellow phenotype for all three gRNAs (Figure 1E). Amp-seq revealed average editing efficiencies of 18.6%, 14.7%, and 9.8%, respectively, much higher (up to 9.8-fold) than wild type (WT) Ymu1 (Weiss et al. 2025) (Figure 1F).

To assess the impact of the WFR variant on multiplexed editing efficiency, we replaced the WT Ymu1 sequence in pTW2278, pTW2279, and pTW2498 with Ymu1-WFR (Figure S3B). Following TRV delivery, we observed much more pronounced phenotypic evidence of editing than with WT Ymu1 (Figure 1G compared with Figure 1B). Amp-seq of yellow sectored plants confirmed enhanced editing: pTW2657 and pTW2658 averaged 48.9% for *AtCHLl1* (gRNA4) and 41.3% for *AtPDS3* (gRNA12) (Figure 1H), while pTW2655 averaged 45.3% and 28.1% at the two *AtCHLl1* sites (Figure 1I). Consistent with the protoplast result (Figure S2H), PCR analysis using primers spanning the *AtCHLl1* gRNA4 and gRNA6 sites revealed large deletions (Figures S3C, S3D).

To characterize germline transmission of multiplexed *AtPDS3* gRNA12 and *AtCHLl1* gRNA4 edits, we selected two plants infected with WT Ymu1 (pTW2278_17 and pTW2278_54) and two plants infected with Ymu1-WFR (pTW2658_5 and pTW2658_80). We observed yellow progeny at frequencies of 4.7% and 1.7% using WT Ymu1, and 6.1% and 12.9% with Ymu1-WFR (Figure 1J, S3E, Table S1). Among the yellow seedlings, the majority of them harbored biallelic edits at both loci (Figure 1J, S3F). Additionally, targeting *AtCHLl1* with two gRNAs, gRNA4 and gRNA6 (using pTW2655), resulted in 8.2% and 24.1% yellow seedlings, with 36/158 (22.8%) of the progeny harboring homozygous large deletions between the two targets (Figures 1J, S3G-I). These data demonstrate that TRV effectively delivers Ymu1-WFR and multiple gRNAs for efficient multiplexed germline editing, that biallelic editing at one locus is highly predictive of biallelic editing at the second target site, and that co-targeting the same gene gave progeny with large deletions between the two target sites.

By optimizing the gRNA array design, and incorporating the highly active engineered Ymu1-WFR variant, we developed an efficient multiplexed editing platform that bypasses the need for transgenesis. Given the broad host range of TRV, we anticipate this approach will be adaptable to many crop species, for example tomato where germline editing has already been demonstrated (Liu et al. 2025). Additionally, the ability to generate large deletions should expand this system’s utility for regulatory element engineering. Finally, this multiplexed system may enable the study of embryonic lethal genes by utilizing *AtCHLl1* as a visual marker; the yellow somatic sectors should facilitate the identification of tissue harboring biallelic knockout of a gene of interest.

## Acknowledgements

Supported by NSF PGRP grant (2334027) to S.E.J. and J.A.D., and Jane Coffin Childs Memorial Fund for Medical Research and NIH (K99GM160778) to H.S. S.E.J and J.A.D. are HHMI Investigators. We apologize to colleagues whose work we could not reference due to space limitations.

## Author Contributions

T.W. and S.E.J. designed research. T.W., H.S., J.A.D. and S.E.J. interpreted data. T.W. and S.E.J. wrote the paper. T.W., M.K., H.S., G.W., A.I., J.A., K.V., M.T., Z.L., E.F., N.S., C.A., and K.C. performed experiments.

## Data Availability

Amp-seq data is accessible at NCBI Sequence Read Archive BioProject PRJNA1427739. RNA-seq data is available at GEO accession: GSE316183.

## Supplementary Materials

**Supplementary Figure 1:**
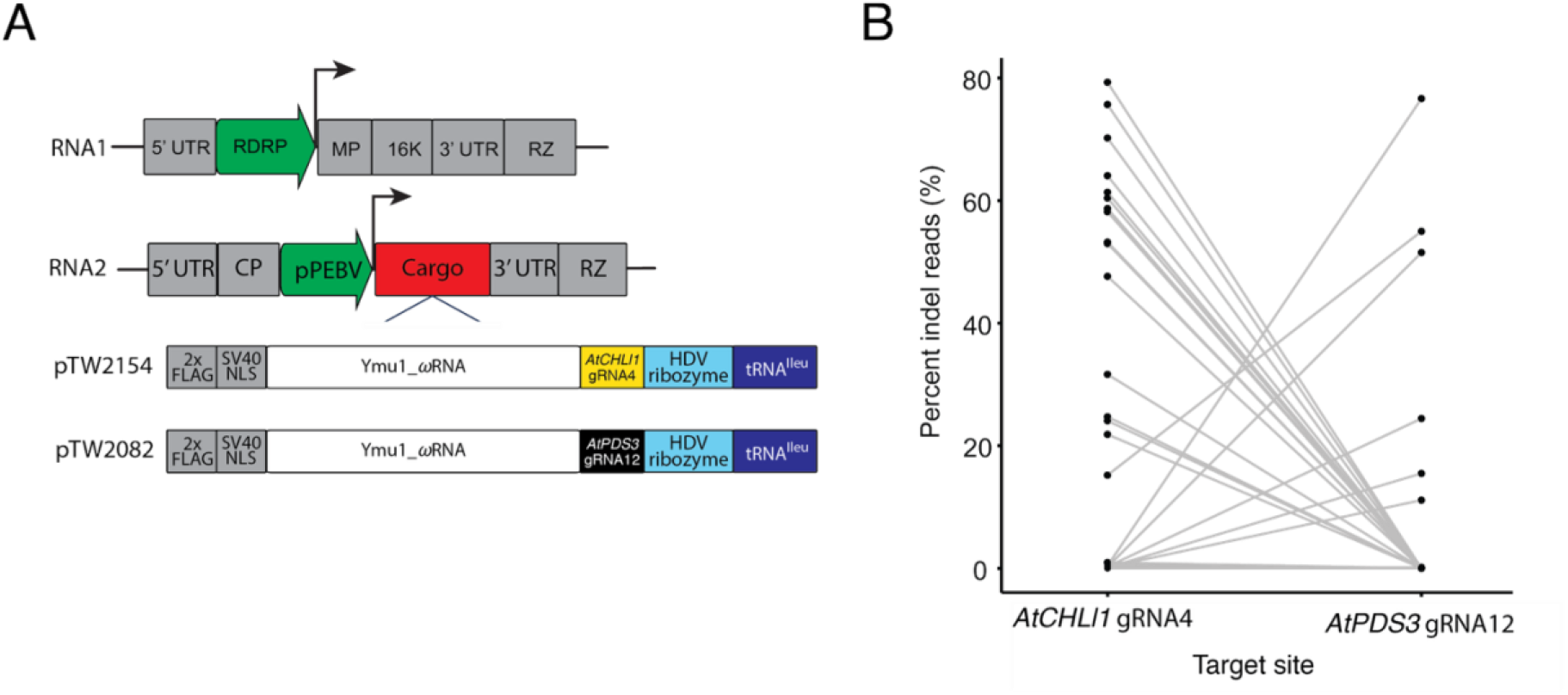
Co-delivery of RNA1 and two RNA2 vectors does not enable multiplexed genome editing. **(A)** Schematic representation of TRV RNA1 and RNA2. The cargo and plasmid ID are listed below RNA2. **(B)** Editing efficiency (y-axis) of plants co-infected with RNA1 and two RNA2 vectors targeting *AtCHLl1* gRNA4 and *AtPDS3* gRNA12 (x-axis). Each gray line represents a single plant, with the black dot at each end of the gray line corresponding to the editing efficiency at each target site.

**Supplementary Figure 2:**
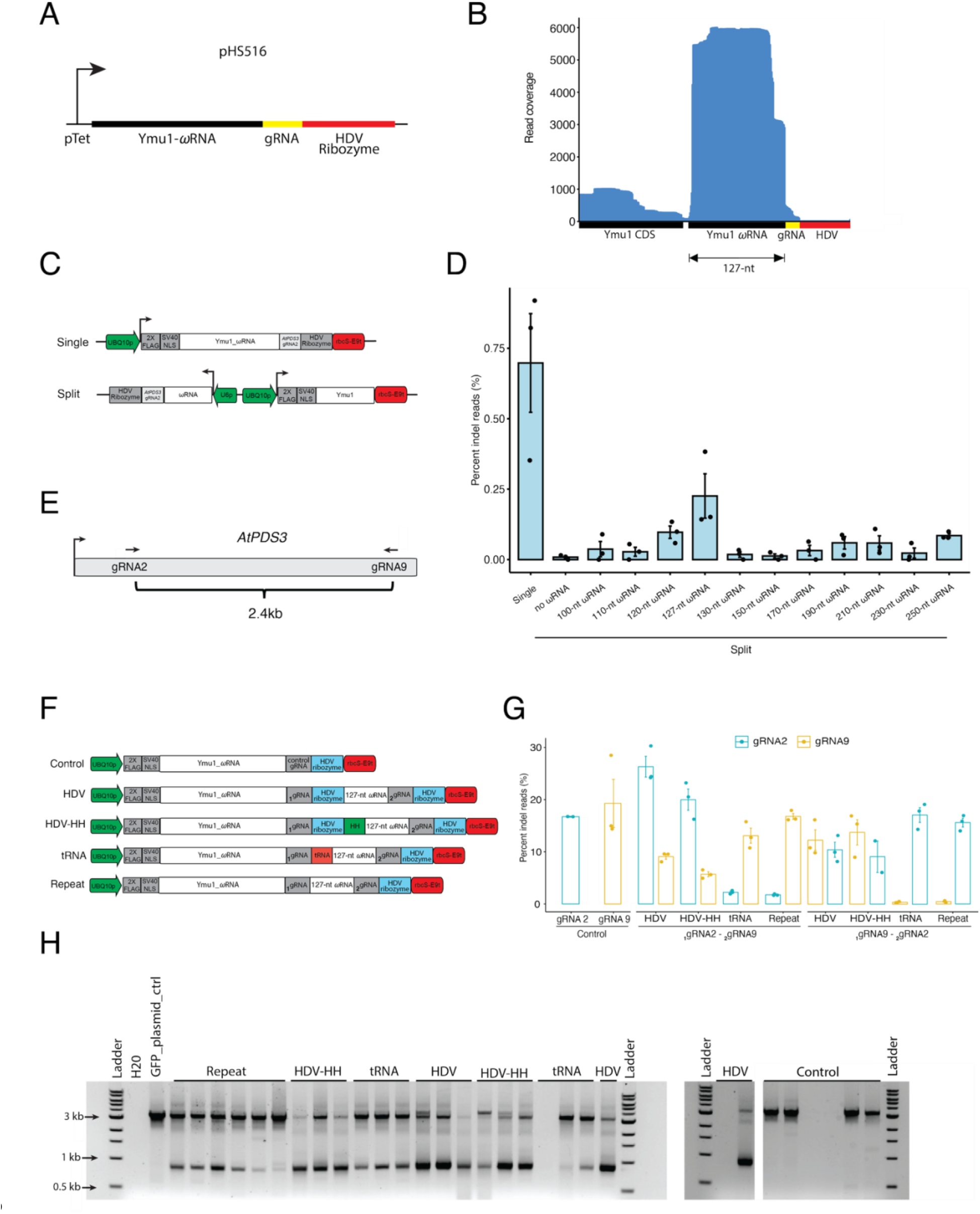
Development of Ymu1 multiplexed gRNA arrays for plant gene editing. **(A)** Schematic of plasmid used in *E. coli* RNA-seq experiment. The tetracycline promoter (pTet, arrow) was used to express the Ymu1-ωRNA (black), gRNA (yellow), and HDV ribozyme (red). The plasmid name (pHS516) is listed above the schematic. **(B)** RNA-seq in *E. coli* using the plasmid shown in panel A. Read coverage (y-axis) is displayed for each Ymu1 single transcript feature (x-axis). **(C)** Schematic of plasmids expressing the Ymu1_ωRNA in a single transcript expression format or split expression format targeting *AtPDS3* gRNA2. In the split expression plasmids the Ymu1 coding sequence was expressed by the *AtUBQ10* promoter (UBQ10p) and the ωRNA was expressed using the U6 promoter (U6p). **(D)** *AtPDS3* gRNA2 editing efficiency (y-axis) using single expression and split expression constructs with varying lengths of ωRNA (x-axis) as shown in panel C. Each dot represents an individual transfection with bars indicating the standard error of the mean (SEM). **(E)** Schematic of *AtPDS3* gene with the arrows indicating the orientation of gRNA2 and gRNA9, located roughly 2.4kb apart. **(F)** Schematic of vectors used in the *Arabidopsis* protoplast multiplexed editing experiment. **(G)** Editing efficiency (y-axis) of the multiplex arrays (x-axis) tested in *Arabidopsis* protoplast targeting *AtPDS3* (gRNA2 and gRNA9). The subscript below each gRNA indicates its placement in the multiplex array as depicted in panel F. **(H)** PCR gel electrophoresis image using primers spanning *AtPDS3* gRNA2 and *AtPDS3* gRNA9 target sites. Deletion between the two target sites corresponds to a ∼850bp band.

**Supplementary Figure 3:**
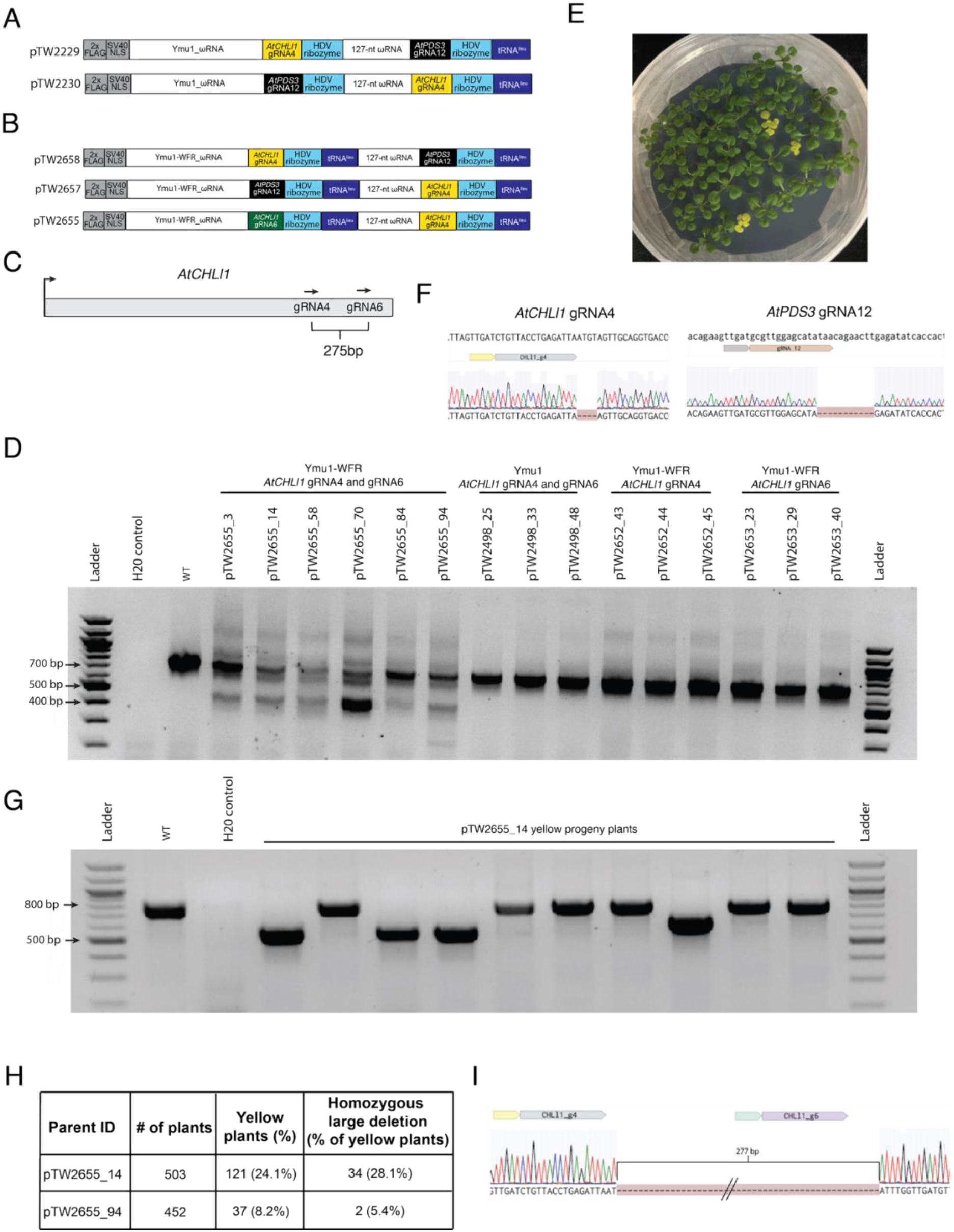
TRV-delivery of Ymu1 for multiplexed genome editing. **(A)** Schematic representation of the multiplexed cargo being expressed by the PEBV promoter in TRV RNA2 targeting *AtCHLl1* gRNA4 and *AtPDS3* gRNA12, containing a single 3’ tRNA^Ileu^. The plasmid ID is listed to the left of each construct. (**B**) Schematic representation of the Ymu1-WFR multiplexed gRNA cargo being expressed by the PEBV promoter in TRV RNA2. *AtCHLl1* gRNA4 (yellow), *AtPDS3 gRNA12* (black), and *AtCHLl1* gRNA6 (green) were used in the multiplex TRV experiments. The plasmid ID is listed to the left of each construct. (**C**) Schematic of *AtCHLl1* gene with the arrows indicating the orientation of gRNA4 and gRNA6, located approximately 275 bp apart. **(D)** Somatic editing analysis of infected plants. PCR gel electrophoresis image using primers spanning *AtCHLl1* gRNA4 and *AtCHLl1* gRNA6 target sites. Deletion between the two target sites corresponds to a ∼420bp band. The plant ID (construct_plant) is listed above each well. The TnpB (Ymu1 or Ymu1-WFR) and gRNA(s) is listed above the plant ID. (**E**) Representative image of yellow and green progeny seedlings from plant pTW2278_17. All yellow seedlings harbored biallelic edits at the *AtCHLl1* gRNA4 target site. (**F**) Sanger sequencing trace file screenshots from a yellow seedling in panel E, harboring homozygous edited alleles at *AtCHLl1* gRNA4 (4bp deletion) and *AtPDS3* gRNA12 (11bp deletion). (**G**) Representative PCR gel electrophoresis image of progeny from pTW2655_14 using primers spanning *AtCHLl1* gRNA4 and gRNA6 target sites. Image shows the presence of deletion between *AtCHLl1* gRNA4 and gRNA6. (**H**) Heritability data from plants pTW2655_14, and pTW2655_94 infected with TRV targeting *AtCHLl1* gRNA4 and gRNA6 (pTW2655). Large deletions were classified as greater than 275 bp deletion between the two target sites. (**I**) Sanger sequencing trace file screenshot from a yellow plant targeted with *AtCHLl1* gRNA4 and gRNA6 harboring a homozygous large deletion (277 bp).

**Supplementary Table 1:** Genotype characterization of green progeny from plants infected with TRV expressing Ymu1 and gRNAs targeting *AtCHLl1* gRNA4 and *AtPDS3* gRNA12 (pTW2278_54 and pTW2278_17).

| Parent ID | pTW2278_54 (# of progeny) | pTW2278_17 (# of progeny) |
| --- | --- | --- |
| # of green plants sequenced | 191 | 232 |
| WT/- *AtCHLI1*,<br>-/- *AtPDS3* | 8 | 11 |
| WT/WT *AtCHLI1*,<br>-/- *AtPDS3* | 9 | 20 |
| WT/- *AtCHLI1*,<br>WT/- *AtPDS3* | 2 | 5 |
| WT/- *AtCHLI1*,<br>WT/WT *AtPDS3* | 2 | 4 |
| WT/WT *AtCHLI1*,<br>WT/- *AtPDS3* | 6 | 33 |
| WT/WT *AtCHLI1*,<br>WT/WT *AtPDS3* | 164 | 159 |

**Supplementary Table 2:**
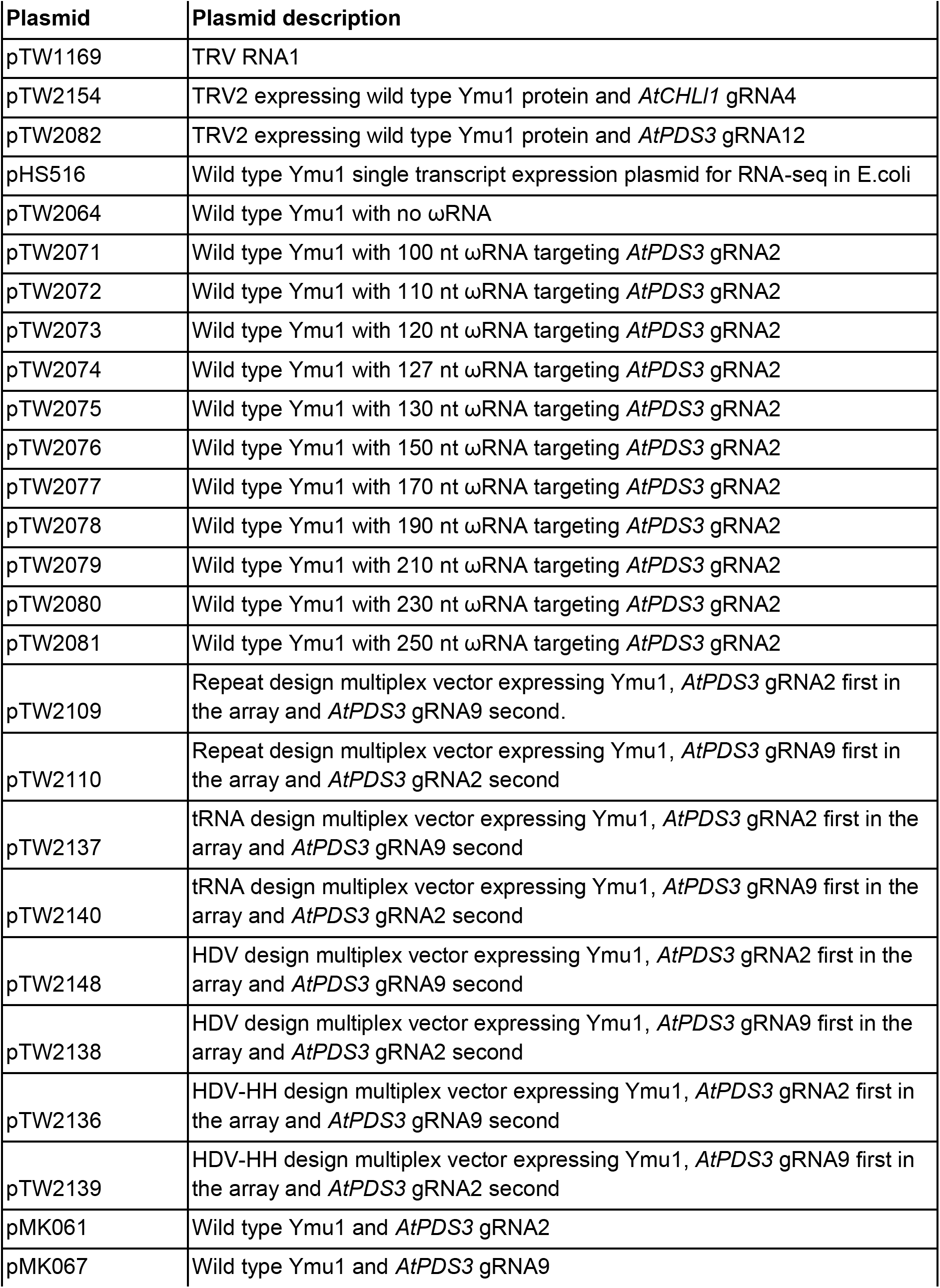
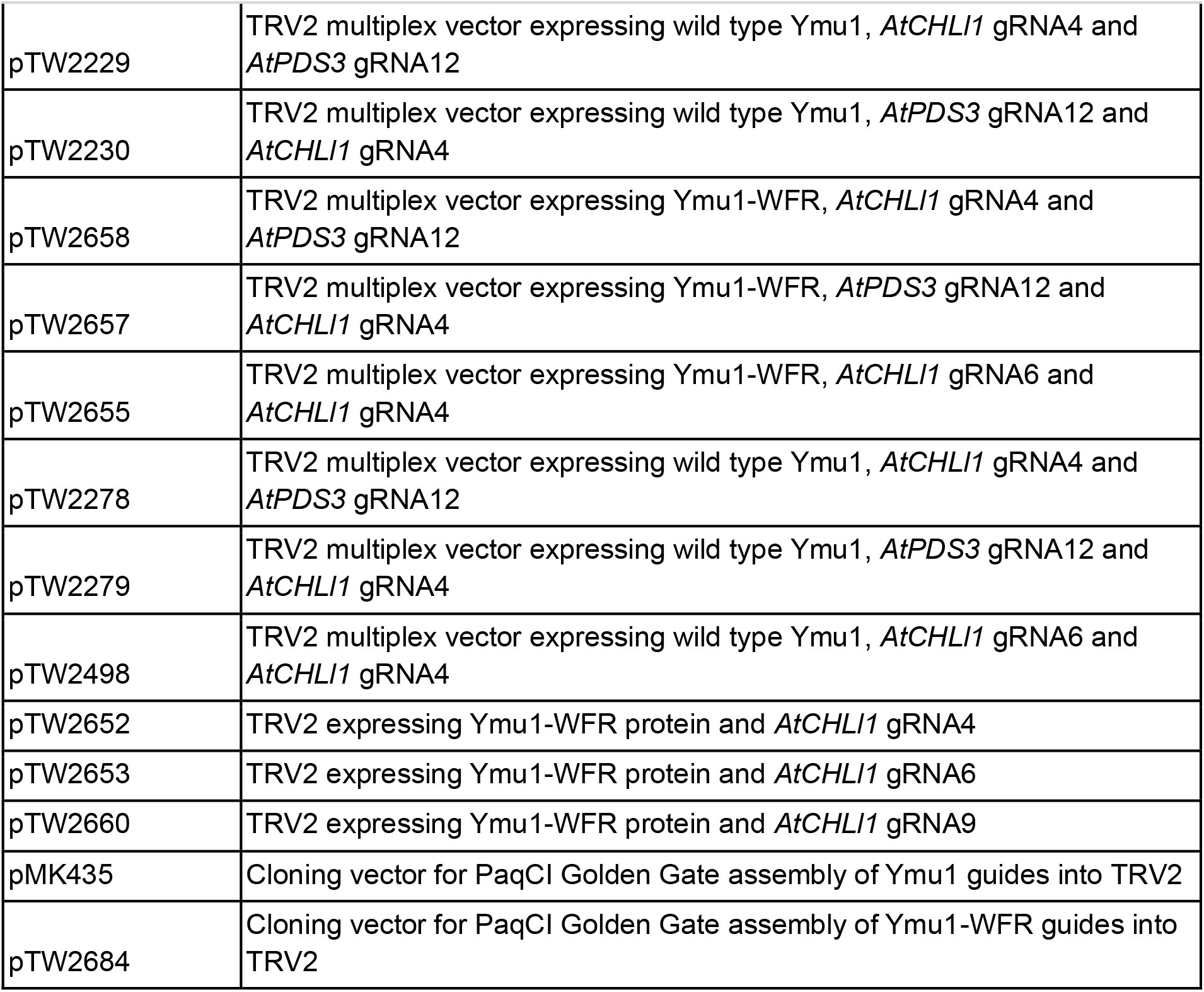
Plasmids and their descriptions used in this study.

**Supplementary Table 3:**
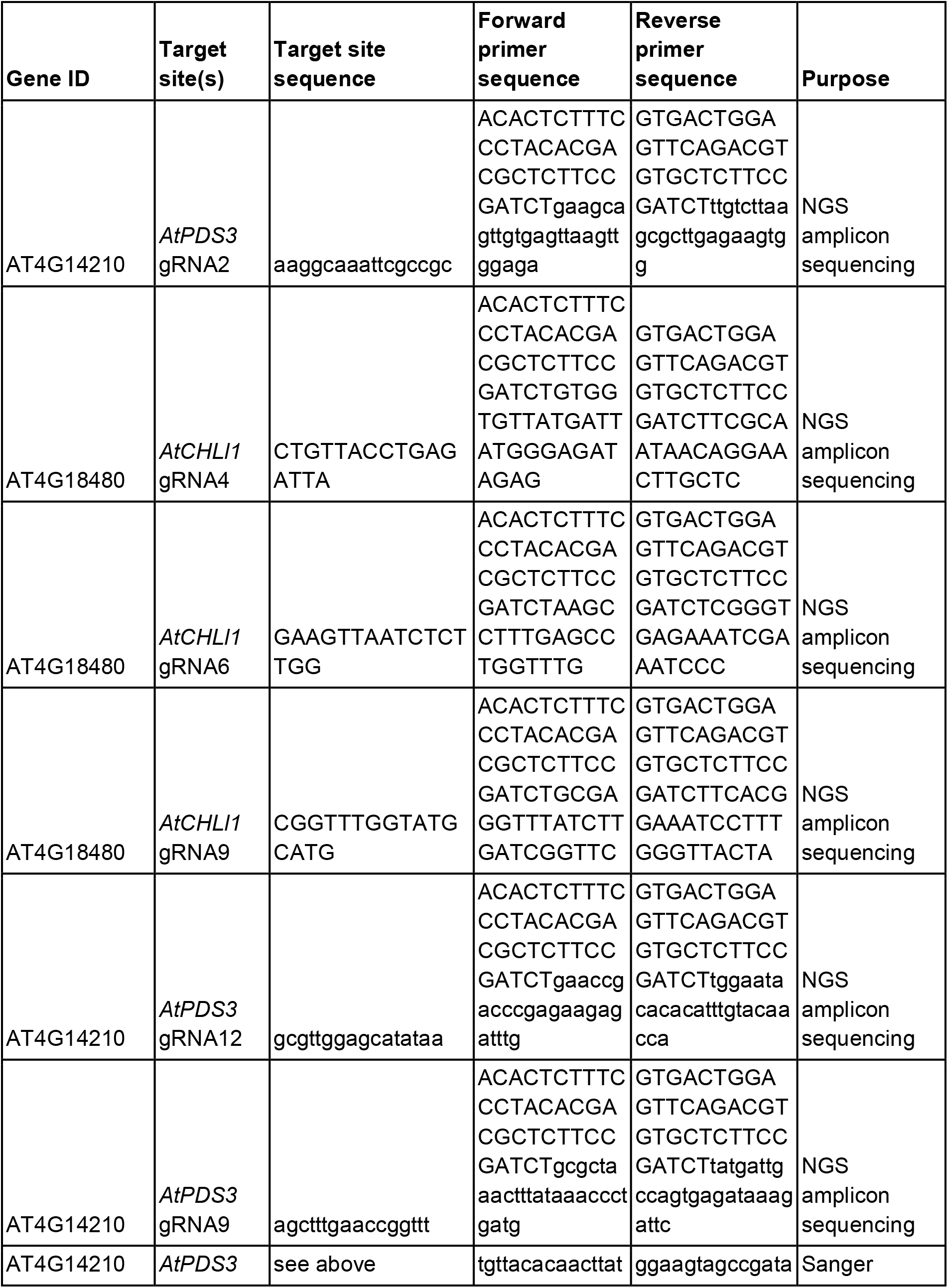
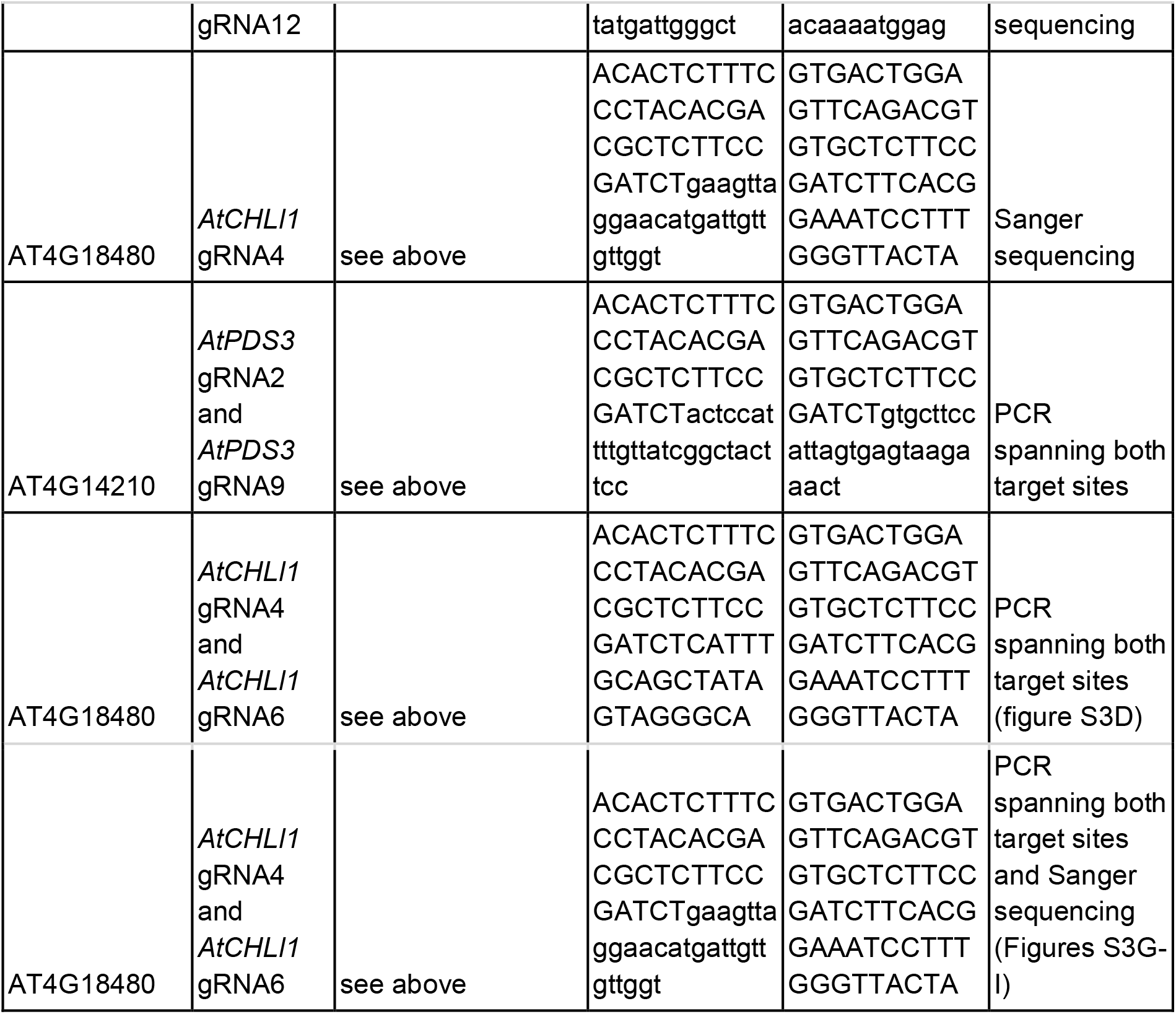
Target sites and primers used in this study.

## Supplementary Methods

### Plasmids used in this study

To generate the Ymu1 plasmids with varying ωRNA lengths, NEB HiFi assembly was performed using two PCR amplicons and pTW2064 (the no ωRNA plasmid backbone). One amplicon contained the U6 promoter and the other amplicon contained the ωRNA. First, pTW2064 underwent a restriction enzyme digestion with SpeI (R3133) and Quick CIP (M0525) to linearize the plasmid. Next, both amplicons and the digested and purified pTW2064 plasmid were used in an NEBuilder HiFi DNA Assembly (E2621) reaction.

Multiplex vectors used in the protoplast experiment expressing wild type Ymu1, *AtPDS3* gRNA2 and *AtPDS3* gRNA9 were created using a golden gate reaction. PCR amplicons or gene fragments (Twist Bioscience) containing the first gRNA, processing sequence, 127-nt ωRNA and the second gRNA were inserted into the pMK025 vector (Weiss et al. 2025) using a PaqCI (R0745) golden gate reaction.

Multiplex TRV2 vectors were created using a golden gate reaction assembly. First, PCR was performed to generate two amplicons; one amplicon contained the first gRNA in the array, an HDV ribozyme and tRNA^Ileu^, and the second amplicon contained the 127-nt ωRNA and second gRNA in the array. Amplicons contained overhangs with PaqCI (R0745) sites to facilitate assembly. Both amplicons and pMK435 (Weiss et al. 2025) were then used in a PaqCI golden gate reaction.

TRV2 vectors containing the Ymu1-WFR variant were created using NEBuilder HiFi DNA Assembly (E2621). The TRV2 vectors were constructed by digesting the wild type Ymu1 TRV2 vectors with HpaI (R0105) and Eco53KI (R0116) restriction enzymes. A gene fragment (Twist Bioscience) containing the Ymu1-WFR sequence was then cloned into the digested plasmid using NEBuilder HiFi DNA Assembly (E2621).

Following assembly, all plasmids were transformed into NEB 10-beta (C3019) *E*.*coli* and validated by whole-plasmid sequencing. Plasmid names and descriptions can be found in Table S2. To enable widespread use of single-target and multiplexed TRV2 vectors containing the Ymu1-WFR variant, we developed a TRV2 ccdb guide dropout vector (pTW2684) and a cloning protocol (Supplementary Protocol).

### Plant gene editing experiments

*Arabidopsis* mesophyll protoplasts were isolated according to previously described methods (Yoo et al. 2007). For protoplast transfection, 20 µg of plasmid was placed at the bottom of each tube and brought to a total volume of 20 µl with water. Then, 200 µl of protoplasts were added to the plasmid solution, followed by 220 µl of fresh, sterile polyethylene glycol (PEG)-CaCl_2_solution. The samples were mixed by gentle tapping and incubated at room temperature for 10 minutes. Transfection was terminated by adding 880 µl of W5 solution and inverting the tubes two to three times. Protoplasts were subsequently pelleted by centrifugation at 100 RCF for 3 minutes, resuspended in 1 ml of WI solution, and plated in 6-well plates precoated with 5% calf serum. The cells were incubated at 26°C for 48 hours. During this incubation period, a 37°C heat-shock treatment was applied for 2 hours, beginning 16 hours post-transfection. At 48 hours post-transfection, the protoplasts were collected for genomic DNA extraction.

TRV delivery was performed as previously described (Weiss et al. 2025; Nagalakshmi et al. 2022). TRV1 and TRV2 vectors were introduced into *Agrobacterium tumefaciens* strain GV3101, and cultures were grown in 200 ml of lysogeny broth (LB) with antibiotics for 18 hours at 28°C. The cultures were centrifuged at 3,500 × g for 20 minutes; the supernatant was discarded, and the pellets were resuspended in 200 ml of sterile water. This suspension was centrifuged again at 2,109 × g for 10 minutes. After discarding the supernatant, the pellet was resuspended in sterile agro-infiltration buffer (10 mM MgCl_2_, 10 mM MES, and 250 µM acetosyringone) to a final optical density OD_600_ of 1.5 and incubated at 23°C for 3 hours with slow shaking. Following incubation, the *Agrobacterium* cultures harboring TRV1 and TRV2 were mixed in a 1:1 ratio (co-delivery experiment used a ratio of two parts TRV1, one part pTW2154, and one part pTW2082). 15 ml of this mixture was delivered to 10-day-old seedlings via agroflood co-culture. After 4 days, the seedlings were transplanted to soil and grown at room temperature for the remainder of their life cycle.

Approximately 12 weeks after viral delivery, seeds were harvested from TRV-infected plants. Seeds were sown on ½ MS plates supplemented with 3% sucrose and stratified at 4°C in the dark for 5 days. Following stratification, the seeds were moved to a growth room under a 16-h light/8-h dark cycle at 23°C for 10–12 days. A subset of plants was then sampled for genotyping by collecting a single piece of leaf tissue. DNA was extracted using the Invitrogen Platinum Direct PCR Universal Master Mix (A44647500) according to the manufacturer’s instructions. Finally, the extracted DNA was analyzed via PCR and Sanger sequencing using the primers listed in Table S3.

### Amplicon sequencing and analysis

Genomic DNA was extracted from protoplast samples using the Qiagen DNeasy Plant Mini Kit (Qiagen, 69106). For plants subjected to agroflood, three leaf tissue samples distal to the TRV delivery site were pooled per plant about 3-4 weeks after viral delivery. The ratio of green to yellow tissue samples collected were selected in a manner that most accurately reflected the phenotype of the entire plant. Once collected, all tissue samples were frozen overnight at −80°C and ground. DNA was subsequently extracted using the Invitrogen Platinum Direct PCR Universal Master Mix (A44647500) according to the manufacturer’s instructions. The DNA was then used for next-generation amplicon sequencing.

Amplicon sequencing was performed using single-end sequencing on the Illumina NovaSeqX platform. Libraries were constructed using a two-step PCR approach. First, target regions were amplified for 25 cycles using primers flanking the TnpB target site (Table S3), followed by purification with 1.0× AMPure XP beads (Beckman Coulter, A63881). Then, samples underwent 12 additional cycles of amplification to attach Illumina indexing primers, followed by a 0.7× AMPure XP cleanup. Final libraries were normalized and pooled for sequencing. Amplicon sequencing analysis was performed using the CrispRvariants R package (v.1.14.0) as previously described (Weiss et al. 2025).

### Bacterial RNA expression and sequencing

For heterologous RNA expression, *E. coli* transformed with pHS516 (Figure S2A, Table S2) and grown in LB medium to an OD_600_ of 0.6 and induced with 10 mM arabinose at 16°C overnight. Following overnight growth, total RNA from cells was extracted using a hot formamide method, in which pelleted *E. coli* were resuspended in an 18 mM EDTA and 95% formamide solution and lysed by incubating at 65°C for 5 minutes. After centrifugation to pellet cell debris, the supernatant containing total RNA was purified using the RNA Clean & Concentrator-5 Kit (Zymo Research R1013) following manufacturer protocols. Approximately 200 ng of purified total RNA was subjected to rRNA depletion using the NEBNext rRNA Depletion Kit for Bacteria (New England Biolabs E7850L) following manufacturer protocols. rRNA-depleted samples were again purified using the RNA Clean & Concentrator-5 Kit and subjected to end repair and small-RNA-seq library prep as previously described (Zhou et al. 2026). Sequencing was performed using an Illumina NextSeq 1000/2000 P2 v3 kit. Following sequencing, paired-end reads were trimmed and merged using fastp (v0.23.2) (Chen et al. 2018). Merged reads of 100–150 nucleotides were selected with fastq-filter (v0.3.0; https://github.com/LUMC/fastq-filter) and aligned to the Ymu1 locus using BWA (v0.7.17) (Li 2013). Alignments were sorted with SAMtools (v1.17) (Li et al. 2009) and converted to BigWig format using deepTools2 (v3.5.1) (Ramírez et al. 2014), then visualized in the Integrative Genomics Viewer (IGV) (Robinson et al. 2011).

## Supplementary Protocol

### Protocol for cloning guides into pTW2684

pTW2684 is a TRV2 vector containing Ymu1-WFR and the ωRNA driven by the PEBV sub genomic promoter. In place of the 16bp gRNA spacer there is a ∼1700 bp insert carrying the toxic ccdb gene and chloramphenicol (CAM) resistance. The insert is flanked by PaqCI Type-II restriction sites, enabling golden gate cloning to replace the toxic ccdb insert with the desired guide sequence.

#### Single target site

1) Order the 16bp guide sequence with 4 bp overhangs as complimentary single stranded oligos from IDT and anneal and phosphorylate them as described below: Top Oligo: 5’-<u>TCAA</u>XXXXXXXXXXXXXXXX-3’ Bottom Oligo: 3’-YYYYYYYYYYYYYYYY<u>CCGG</u>-5’ *Underlined nucleotides are complimentary to the overhangs created by PaqCI digestion of pTW2684 Phosphorylation and annealing reaction setup: 3 μL 100 μM Top oligo 3 μL 100 μM Bottom oligo 3 μL T4 DNA ligase buffer (with ATP) 2 μL T4 polynucleotide kinase (PNK) 19 μL Water
2) Place samples in a thermal cycler and run the following program: 37°C/30 min + 95°C/5 min + 90°C/1 min + 85°C/1 min + 80°C/1 min + 75°C/1 min + 70°C/1 min + 65°C/1 min + 60°C/1 min + 55°C/1 min + 50°C/1 min + 45°C/1 min + 40°C/1 min + 35°C/1 min + 30°C/1 min + 25°C/hold
3) Next, set up a golden gate reaction to clone the annealed oligos into pTW2684. Golden gate reaction setup: 50 ng pTW2684 μL annealed oligos (diluted 1:25) 0.5 μL PaqCI enzyme 0.5 μL PaqCI Activator 2 μL T4 DNA ligase buffer 1 μL of T4 DNA ligase Water up to 20 μL
4) Place samples in a thermal cycler and run the following program: 10x(37°C/5min + 16°C/10min) + 37°C/15min + 80°C/5min + 4°C hold
5) Transform 2-5 μL of the reaction into NEB 10-beta (C3019) *E*.*coli* cells and select for successfully cloned plasmids using kanamycin plates.

#### Multiplexed vector containing two target sites

Cloning two gRNAs into pTW2684 requires a golden gate reaction with two PCR amplicons. This example describes the cloning strategy to build pTW2658 (*AtCHLl1* gRNA4 and *AtPDS3* gRNA12). To clone different gRNAs you can either first clone each guide separately into pTW2684 using the protocol above, and then use that plasmid as a PCR template, or order synthesized DNA fragments.

1) Perform two PCR reactions and run the samples on a gel to confirm the correct size. (rxn 1 is 192 bp and rxn 2 is 185 bp). Purify the PCR reaction and nanodrop.

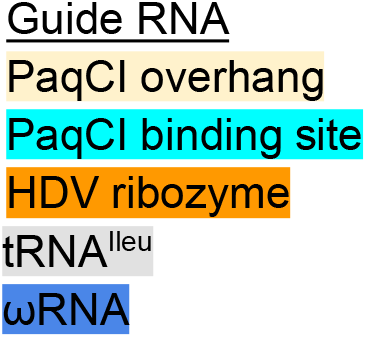

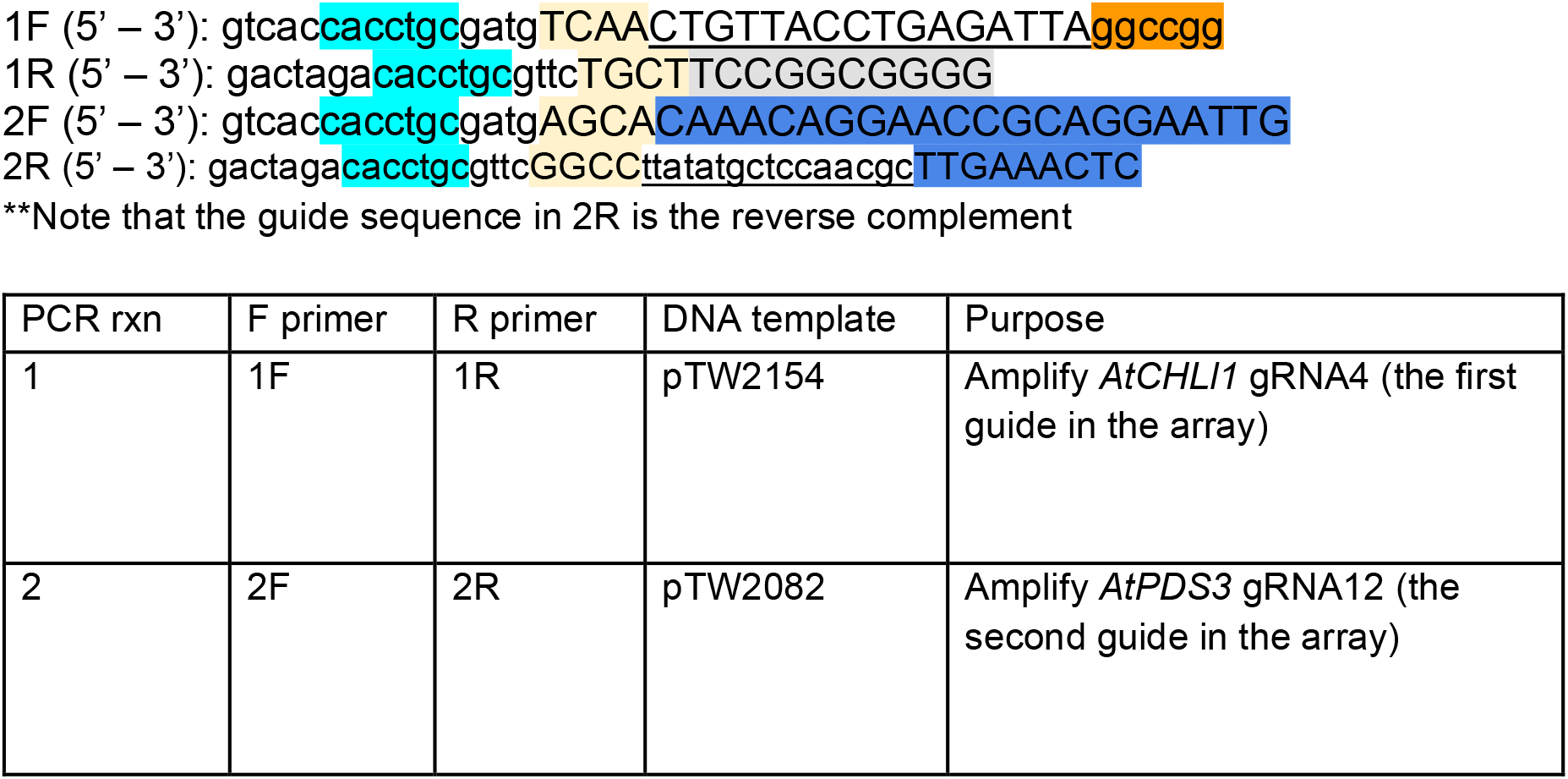

#### PCR reaction

2.5 µl F primer 10 μM

2.5 µl R primer 10 μM

2 ng Template plasmid DNA

25 µl Q5 Hi Fidelity DNA polymerase master mix

Water up to 50 µl

Place samples in a thermal cycler and run using the following settings:

1. 98°C - 30 sec
2. 98°C - 10 sec
3. 55°C - 20 sec
4. 72°C - 20 sec
5. Repeat steps 2-4 35x
6. 72°C - 2 min
7. 4°C - hold

#### Amplicon 1

Sequence (5’ – 3’):

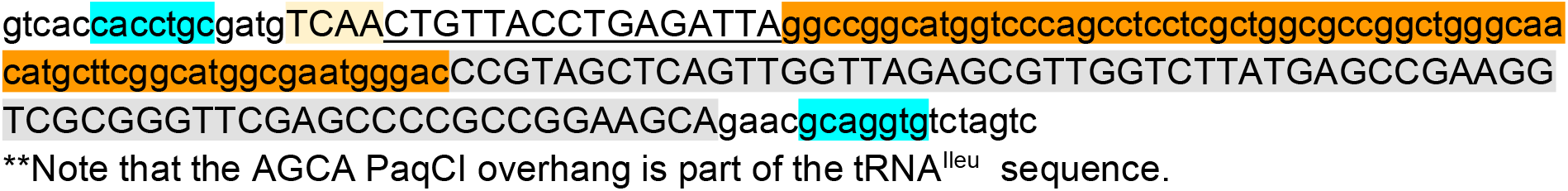

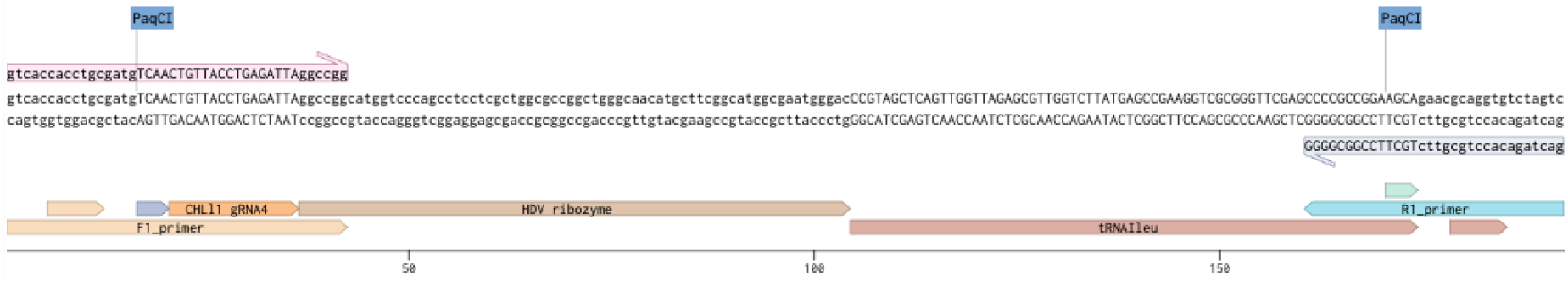

#### Amplicon 2

Sequence (5’ – 3’):

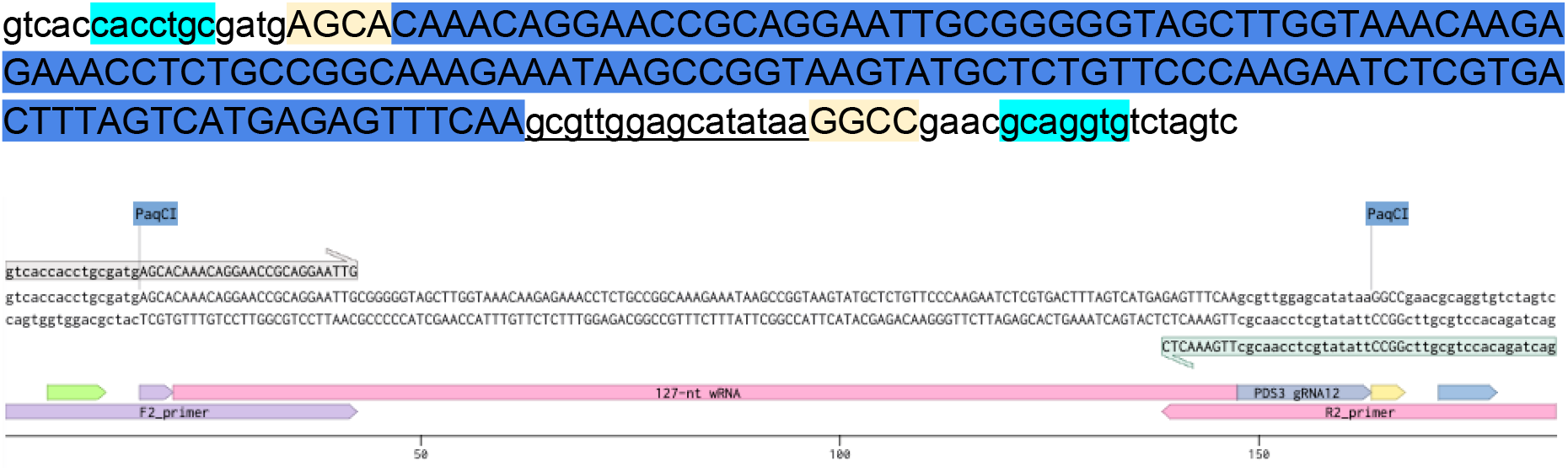

(2) Perform golden gate reaction 100 ng pTW2684 100 ng amplicon 1 100 ng amplicon 2 0.5 µl PaqCI 0.5 µl PaqCI activator 1 µl T4 DNA ligase 2 µl T4 DNA ligase buffer Water up to 20 µl

Place samples in thermal cycler and run the following program: 10x(37°C/5min + 16°C/10min) + 37°C/15min + 80°C/5min + 4°C hold

(3) Transform 2-5 μL of the reaction into NEB 10-beta (C3019) *E*.*coli* cells and select for successfully cloned plasmids using kanamycin (KAN) plates.

